# Song sparrows do not discriminate between their own song and stranger song

**DOI:** 10.1101/2020.05.08.084665

**Authors:** Çağlar Akçay, Michael D. Beecher

## Abstract

Bird song is socially learned. During song learning, the bird’s hearing its own vocalization is important for normal development of song. Whether bird’s own song is represented and recognized as a special category in adult birds, however, is unclear. If birds respond differently to their own songs when these are played back to them, this would be evidence for auditory self-recognition. To test this possibility, we presented song sparrow males (*Melospiza melodia*) playbacks of their own songs or stranger songs and measured aggressive responses as well as type matching. We find no evidence of behavioral discrimination of bird’s own song relative to the (non-matching) stranger song. These findings cast doubt on an earlier proposal that song sparrows display auditory self-recognition and support the common assumption in playback experiments that bird’s own song is perceived as stranger song.

## Introduction

Songbirds are one of the handful of taxa (including humans, bats, cetaceans, parrots and hummingbirds) that are known to learn their vocalizations socially. Laboratory studies of songbirds have revealed that a young bird’s hearing its own vocalizations plays a key role in song development: young birds deprived of auditory feedback from their own singing ultimately develop abnormal songs (Brainard & Doupe, 2000; Konishi, 1965; Price, 1979). Hearing one’s own singing also appears to be important for the maintenance of adult song: birds deafened as adults also lose the species typical consistency in their songs (Lombardino & Nottebohm, 2000; Nordeen & Nordeen, 1992). Not surprisingly, song-learning and production circuits in the songbird brain display special sensitivity to own song when these are played back from a speaker (Poirier, Boumans, Verhoye, Balthazart, & Van der Linden, 2009; Prather & Mooney, 2004). All these results suggest that songbirds may represent their own song as a special category and may even display auditory self-recognition (Derégnaucourt & Bovet, 2016).

Field studies have raised different questions as to ways in which a bird’s own song might be special. Bird song, which typically functions in territory-defense and mate-attraction contexts (Catchpole & Slater, 2008), is often learned from other adults in the local area, usually a neighbour (Beecher, 2008). Consequently, in natural interactions, birds will often hear versions of their songs from other individuals. Bird song communication is often studied by observing a bird’s response to the playback of pre-recorded song (McGregor, 2000; Weeden & Falls, 1959). Many such playback experiments simulate realistic situations, such as an intrusion onto the subject’s territory by a neighbour, or a stranger but a particularly interesting case is when the bird’s own song is played back to him. Playback of own song of course would not simulate a realistic situation if the bird were to perceive the playback as its own song. It may however resemble a realistic situation if the bird perceives the playback song as the song of a stranger that happens to have a song very similar to his, or as a shared song of one of his neighbours. The neighbour possibility is plausible in those populations where neighbours share songs, a common occurrence for species who learn songs after they disperse from the natal area, such as indigo buntings, *Passerina cyanea*, or some populations of song sparrows, *Melospiza melodia*, (Beecher, Campbell, & Stoddard, 1994; Payne, 1982). The stranger possibility is plausible in circumstances where some birds learn songs in one neighbourhood and then move to another neighbourhood leading to shared songs between distant birds. The self-recognition possibility seems least likely although McArthur (1986, 1987) claimed to have found evidence for this possibility in a study with song sparrows (see below).

One difficulty in studying auditory self-recognition in the field is that it is unclear how self-recognition would manifest itself in the response of an individual to playback (Derégnaucourt & Bovet, 2016). The first possibility is that the bird might type match (i.e. reply with the same song type as the playback song) its own song at a higher rate than it would to, say, a neighbour song, as has been found in western meadowlarks, *Sturnella neglecta* (Falls, 1985). Note that such an effect could simply come about because bird’s own song is by definition more similar (indeed identical) to his than any other class of stimulus (Falls, Krebs, & McGregor, 1982). In any case, other studies found different patterns. For instance, Stoddard and colleagues (1992) found that type matching levels in male song sparrows were similar to own song and to matching stranger songs, while type matching in response to matching neighbor songs were lower than either. Since stranger matching songs were generally worse matches to subject songs than either own- or neighbour matching songs, similarity of playback stimulus to the subject song cannot explain the pattern of matching found in this experiment.

Other studies assessed whether birds differ in aggressive responses to own song vs. stranger or neighbour song (Derégnaucourt & Bovet, 2016). Several studies have found an intermediate response to the own song, i.e., stronger than to neighbour song, and weaker than to stranger song (Brooks & Falls, 1975; Weeden & Falls, 1959; Yasukawa, Bick, Wagman, & Marler, 1982). In contrast, Searcy and colleagues (1981) found no significant difference in approach to own song compared to stranger song in eastern song sparrows. Most relevant to the present study, McArthur (1986) found that song sparrows responded with equal intensity in singing, flights or approach measures to own song compared to stranger song while the playback was ongoing, but birds stayed closer to the speaker after the playback was over in the two stranger song conditions compared to the own song condition. McArthur interpreted this one difference in his six response measures as evidence of ‘auditory concept of self’ (McArthur, 1986). This interpretation has been criticized (Suarez & Gallup, 1987).

In the present study we studied the same western population studied by Stoddard et al. (1992). Male song sparrows have a repertoire of about 9 distinct song types (median: 9, range: 6-13 songs) and in this population they show high levels of song sharing among neighbours (Beecher, Campbell, & Stoddard, 1994; Hill, Campbell, Nordby, Burt, & Beecher, 1999). This song sharing comes about by birds copying whole songs of older birds (‘tutors’), who are often also neighbours (Beecher, Campbell, & Stoddard, 1994; Nordby, Campbell, & Beecher, 1999). This fact means that any particular song in a song sparrow’s repertoire is invariably more similar in acoustic structure to a song in another bird’s repertoire (e.g. the tutors or neighbours) than it is to one of the other songs in his own repertoire. Additionally, song sparrow songs do not have distinctive ‘signature’ or voice characteristics identifying them as the songs of that individual (Beecher, Campbell, & Burt, 1994).

Thus, when we play own song to a bird in the field, it is plausible that he would perceive it as the similar song of one of his neighbours. Song sparrows, like many other territorial animals display reduced aggression towards neighbours compared to strangers, i.e. the Dear Enemy effect (Fisher, 1954; Stoddard, Beecher, Horning, & Campbell, 1991; Temeles, 1994). Thus, if own song is perceived as a neighbour song we expect subjects to respond more weakly to own song than to stranger song (Stoddard et al., 1991). Alternatively, he might perceive own song as from a stranger who happens to share a song with him, given that song types do spread beyond the immediate neighbourhood (Nordby et al., 1999). In this case we would expect him to respond equally strongly to own and stranger song. Finally, it is also possible that playback of own song is especially salient, either because his own songs will have been the songs he has heard more than any others or because it is represented as a distinct category for recognition. In this case, we would expect a difference in response between stranger and own song, although which direction this difference should go is unclear, as discussed above.

We were also motivated to carry out this experiment for methodological reasons: Bird’s own song is a powerful stimulus for studying behaviours like song type matching. Using bird’s own song is also a convenient way to match stimulus song for relative quality, familiarity and similarity to each subject while avoiding pseudoreplication. Consequently, a number of studies have used bird’s own song as a stimulus in playback experiments (e.g. Akçay, Tom, Campbell, & Beecher, 2013; Anderson, Searcy, & Nowicki, 2005; Searcy, Ocampo, & Nowicki, 2019), with the assumption that this song is perceived as a stranger song. The present study provides a test of this assumption.

## Methods

### Study site and subjects

We studied adult male song sparrows in a population in Discovery Park, Seattle, WA that has been studied since 1986 (Beecher, Campbell, & Stoddard, 1994). We carried out this experiment between 17 and 22 June 2019. The subjects used in this study were not banded but we had no more than 4 days between the initial recordings of the songs of subjects and the trials to ensure that the individual recorded for own song playbacks was still there at the time of the trials. Given that territories are extremely stable in song sparrows even across multiple years (e.g. Akçay, Campbell, & Beecher, 2015) this assumption is reasonable. We also corroborated the identities of the birds by matching the song types recorded in the recording sessions before the trials and songs recorded during the trials.

We tested 24 males. The sample size was determined *a priori* with a power analysis, aiming for a power of 0.80, and an effect size of d=0.6. This effect size is smaller than the significant effects we detected in our studies of individual discrimination of different classes of individuals (Akçay, Campbell, & Beecher, 2017; Akçay, Reed, Campbell, Templeton, & Beecher, 2010; Akçay et al., 2009; Stoddard et al., 1991).

### Stimuli and Design

We recorded each subject for at least 15 minutes which was enough to obtain at least two or three good song types. We did not attempt to record the entire repertoire of the males. During these recordings we occasionally used playbacks both to elicit song, and to determine the extent of the territory for the trials. The recordings were made with a directional microphone (Sennheiser ME67/K6) and a digital recorder (Marantz PMD660). We viewed the recordings in Syrinx (John Burt, Portland, OR) and chose one good rendition of a song type for each subject for own song stimulus. We manually filtered low-frequency noise and added a silent period to the end of the song to create a 10 second wave file. This wave file was played during the trial in a loop so that playback stimulus rate was one song every 10 seconds (song sparrow songs last about 3 seconds).

Each stimulus song was used once as an own song and once as a stranger song for a different subject at least 4 territories away (average distance: 667 m, range: 490-961 m). This definition of stranger is consistent with previous studies on this species (Kroodsma, 1976; Searcy et al., 1981; Stoddard et al., 1991). Each subject was tested twice, once with his own song and once with the stranger song in a counterbalanced order such that half of the subjects received the own song first, while the other half received the stranger song first. The trials were carried out on consecutive days with about 24 hrs in between.

The own song stimulus by definition was one which the subject could match. Because we did not attempt to record the entire repertoire of the subjects, we did not ensure that the stranger stimulus song could be matched but we observed no type matching during the stranger trials. Because song sharing with non-neighbours is low in our population (Hill et al., 1999), we assumed that most, if not all of the stranger songs were non-shared with the subject.

### Procedure

For each trial, we placed a Bluetooth speaker (Anker SoundCore, Anker Inc.) on a natural perch at a height of about 1.5m above ground at a central location of the territory. The speaker volume was adjusted to play the stimulus songs at about 85 db SPL measured at 1 m, corresponding to natural levels of broadcast song in this species. We then moved to about 10-15 m from the speaker and started the playback, which lasted for 3 minutes after the subject’s first response (approach or song). This trial duration was the same as in previous studies on recognition of song in song sparrows (Searcy et al., 1981; Stoddard et al., 1991) and initial responses to three minutes of playback predict an eventual attack on a taxidermic mount (Akçay et al., 2013). In all cases, the birds responded within about 10-20 seconds of the start of the playback (to the first or second playback song). During the playback, the observers narrated the trial and recorded the vocal response of the subject using the same equipment as above. We recorded each flight (any airborne movements), distance with each flight, songs (loud and soft, see below), and wing waves. After the playback ended, we recorded the same behaviors for another 2 minutes. We avoided testing neighbours on the same day and carried out consecutive trials at least 200 m apart, where we could not hear the previous subject.

### Response measures

From the trial recordings we extracted the following information: flights of the subject (any airborne movements), distance to the speaker with each flight, loud songs, soft songs and wing waves. Soft songs are low amplitude songs that reliably predict a physical attack (Akçay et al., 2013; Nice, 1943; Searcy, Anderson, & Nowicki, 2006). Loud songs and soft songs can be distinguished in the field by an experienced observer (in this case, CA) despite the fact that amplitude variation in singing is continuous (Anderson, Searcy, Peters, & Nowicki, 2008). We further distinguished between two types of soft songs: crystallized and warbled soft songs which differ in their acoustic structure (Anderson et al., 2008). Wing waves are rapid fluttering of one or both wings without taking off, which also predict a subsequent attack in our population (Akçay et al., 2013). Loud songs, in contrast do not predict attack in this species (Akçay et al., 2013; Searcy et al., 2006). We converted counts of behaviours into rates to account for slight variation in playback duration (mean duration ± SD: 180.66 ± 2.95 seconds). We extracted the same response variables for the 2-minute post-playback duration as well.

### Data analyses

Most of the rates were non-normally distributed. We therefore used permutation tests (1000 iterations) to compare responses to the own song with responses to stranger song. We report the effect sizes and 96% confidence intervals along with the comparisons. Finally, we also report repeatability of the behaviours for completeness sake. The analyses were carried out in R (R Core Team, 2012) using packages ez (Lawrence, 2016) and rptR (Nakagawa & Schielzeth, 2010). To assess whether type matching was greater than random we ran a binomial test. Because we did not record the entire repertoires of the subjects, we did not know their repertoire size to exactly determine the chance level of matching. Previous samples of males in our population have consistently yielded average repertoire sizes larger than eight (e.g. Hill et al., 1999; Nordby et al., 1999). Therefore, we conservatively assumed a chance level of matching of 1/8.

### Ethical Note

The experiments lasted less than 10 minutes between arrival on territory and leaving (with only about three minutes of playback) to minimize disturbance to the subjects. The subjects were not captured at any time for this study. The procedures conform to the guidelines of ASAB/ABS for the treatment of animals in behavioral research and teaching.

## Results

There were no differences between responses to own song and stranger song during either the 3-minutes of playback (Table 1) or the 2-minute period after the playback (Table 2). The response variables during the playback period were highly repeatable (Table 1). Subjects type matched own song playback in 6 of the 24 trials. In an additional 3 trials, the subjects were singing the same song type as playback song when we started (thus we type matched the subject). In those cases, the subjects stayed on type. We counted them as type matching as well. This level of type matching (9/24=0.375) was higher than the expected level of type matching (1/8), p=0.0013. When we excluded the three trials on which the subjects stayed on type the observed type matching levels (6/21=0.286) was still higher than the expected level by a binomial test p=0.028.

**Table 1.** Results of the permutation tests (p-values) for each response variable measured during the trial, along with effect sizes (Cohen’s d and 95% confidence intervals) and repeatability (intra-class correlation coefficients; standard errors are given in parentheses and p-values associated with the coefficients).

| variable | Permutation<br>p-value | Cohen's d | 95% CI | ICC (SE) | ICC p-value |
| --- | --- | --- | --- | --- | --- |
| **Flight rate** | 0.233 | 0.136 | -0.43, 0.71 | 0.86 (0.076) | <0.0001 |
| **Proportion of time spent <1m** | 0.174 | 0.265 | -0.28, 0.90 | 0.58 (0.14) | 0.001 |
| **Warbled soft songs** | 0.17 | 0.277 | -0.25, 0.86 | 0.49 (0.17) | 0.0067 |
| **Crystallized soft songs** | 0.84 | 0.0481 | -0.54, 0.62 | 0.43 (0.18) | 0.01 |
| **Wing waves** | 0.69 | 0.0671 | -0.52, 0.60 | 0.69 (0.12) | <0.0001 |
| **Loud songs** | 0.35 | -0.154 | -0.77, 0.39 | 0.72 (0.095) | <0.0001 |

**Table 2.** Results of the permutation tests (p-values) for each response variable measured during the 2 minutes after the playback, along with effect sizes (Cohen’s d) and their 95% confidence intervals.

| variable | Permutation<br>p-value | Cohen's<br>d | 95% CI |
| --- | --- | --- | --- |
| **Flight rate** | 0.57 | -0.107 | -0.56, 0.24 |
| **Proportion of time spent <1m** | 0.97 | -0.0065 | -0.45, 0.45 |
| **Warbled soft songs** | 0.54 | -0.18 | -0.57, 0.22 |
| **Crystallized soft songs** | 0.43 | 0.17 | -0.30, 0.50 |
| **Wing waves** | 0.64 | 0.22 | -0.12, 0.65 |
| **Loud songs** | 0.19 | 0.266 | -0.13, 0.75 |

## Discussion

We found no evidence that male song sparrows discriminated between their own song and a stranger song: on no response variable were responses to self and stranger songs significantly different. The non-significant differences includes proximity to the loudspeaker after the playback period (as measured by proportion of time spent within 1 m during the 2 min after playback),the only measure of six for which McArthur (1986) found a significant difference between self and stranger conditions, and which served as the basis of his claim for an auditory self-concept. The present results therefore offer no support for the existence of an auditory self-concept in song sparrows and also corroborate an earlier finding of no discrimination between self and stranger songs in song sparrows (Searcy et al., 1981).

The observed levels of type matching in response to own song (9 out of 24 trials) was slightly lower than the matching to own song (6 out of 10 subjects) or to stranger song (5 out of 10 subjects) that was observed in a previous study in our population (Stoddard, Beecher, Horning, & Campbell, 1992) but not significantly so (Fisher’s exact test; p= 0.41and 0.72 respectively). Note also that the song playbacks in that earlier study were carried out in early Spring (March) when type matching interactions are generally higher between neighbouring birds (Beecher, Campbell, Burt, Hill, & Nordby, 2000). Thus, the observed levels of type matching in our present experiment are consistent with previous estimates of song type matching against own or stranger song and corroborates the lack of discrimination between these two categories.

One potential caveat is that lack of discrimination does not always indicate lack of recognition. For instance, song sparrows in our population display the Dear Enemy effect (i.e. reduced aggression towards neighbours compared to strangers) when the songs of these neighbours are played from the correct boundary (Stoddard et al., 1991). When the neighbour and stranger songs are played at the centre or an incorrect boundary however, males respond equally strongly to the neighbour and stranger songs (Stoddard et al., 1991). This is not because of a lack of discrimination, as evidenced by the fact that intrusions by a neighbour are retaliated upon even after the neighbour “retreats” to his own territory (Akçay et al., 2010; Akçay et al., 2009). Rather, the lack of discrimination between strangers and neighbours at the territory centre likely comes about because a bird singing in your own territory is a high threat to territorial integrity or paternity whether he is a stranger or neighbour.

In summary, we think it is unlikely that song sparrows recognize their own song as such, given the only previous evidence for such recognition in song sparrows was based on a very similar field playback test, with similar response measures, and with no statistical correction made for the multiple measures (McArthur, 1986). Indeed, considered together, the four studies in song sparrows (McArthur, 1986; Searcy et al., 1981; Stoddard et al., 1992; and the present study) that measured behavioural responses to own song show little evidence for behavioural discrimination, let alone own-song recognition in this species.

This conclusion is relevant to another recent study in which we found that song sparrows responded more aggressively to their primary tutors song compared to stranger songs, and the difference in response strength is dependent on the amount of song learning from that primary tutor (Akçay et al., 2017). That finding could have come about if the birds somehow perceived the tutor song playback as their own song and somehow responded more aggressively to it. The present results argue against this interpretation as responses to own songs and stranger songs did not differ (despite the fact that the present study has a larger sample size *a priori* chosen to detect a smaller effect than the found in the Akçay et al., 2017).

### Self-recognition in songbirds in other modalities

Whether songbirds other than magpies (Prior, Schwarz, & Güntürkün, 2008) show evidence of self-recognition in any modality is an open question. Multiple recent studies in several species of corvids found that while birds often displayed mirror contingent behaviors they generally do not pass the mirror-mark test (Clary, Stow, Vernouillet, & Kelly, 2020; Kusayama, Bischof, & Watanabe, 2000; Medina, Taylor, Hunt, & Gray, 2011; Soler, Pérez-Contreras, & Peralta-Sánchez, 2014; Vanhooland, Bugnyar, & Massen, 2019). Whether this is due to methodological artifacts or true lack of self-recognition is a matter of some debate (Clary & Kelly, 2016; Clary et al., 2020). Only one other study claimed to have found clear evidence of self-recognition in another corvid; the Indian house crow, *Corvus splendes* (Buniyaadi, Taufique, & Kumar, 2019). Thus, on the whole, the evidence for self-recognition in the visual modality is mixed.

Among non-corvid songbirds, we are aware of only one study that tested self-recognition in the visual modality. In this study great tits, *Parus major*, failed to pass the mirror-test (Kraft, Forštová, Utku Urhan, Exnerová, & Brodin, 2017). Given that many songbirds habitually attack their mirror images (e.g., on the side-view mirrors of cars), as if to confront an intruder (Branch, Kozlovsky, & Pravosudov, 2015; Censky & Ficken, 1982), this result suggests that non-corvid songbirds are unlikely to possess self-recognition.

Finally, as noted above our motivation to carry out this experiment was partly methodological: Studies examining the pattern and signal value of song type matching often face the difficult problem of finding stimulus songs shared by the focal subject while controlling for familiarity and similarity (Falls et al., 1982). Using the bird’s own song, if it is perceived as a stranger song, instantly solves this problem: the own song is matched to each subject in both familiarity and similarity. Several studies have used this approach (e.g. Akçay et al., 2013; Anderson et al., 2005; Searcy et al., 2019). The present study supports the validity of this methodological choice, at least in song sparrows.

## Conclusion

We found no evidence that song sparrows show auditory discrimination of their own song from stranger songs. Instead, own song played back from inside the territory seems to be perceived as a stranger song that is shared or possibly a neighbour intruding. These results validate the use of own song in studies of animal communication.

## Acknowledgements

This study was supported by a Young Investigator Award (BAGEP) to ÇA from the Science Academy of Turkey.

## Notes

### Competing Interest Statement

The authors have declared no competing interest.

## References

Akçay, Ç., Campbell, S. E., & Beecher, M. D. (2015). The fitness consequences of honesty: under-signalers have a survival advantage in song sparrows. Evolution, 69, 3186–3193.

Akçay, Ç., Campbell, S. E., & Beecher, M. D. (2017). Good tutors are not dear enemies in song sparrows. Animal Behaviour, 129, 223–228. doi: https://doi.org/10.1016/j.anbehav.2017.05.026

Akçay, Ç., Reed, V. A., Campbell, S. E., Templeton, C. N., & Beecher, M. D. (2010). Indirect reciprocity: song sparrows distrust aggressive neighbors based on eavesdropping. Animal Behaviour, 80, 1041–1047.

Akçay, Ç., Tom, M. E., Campbell, S. E., & Beecher, M. D. (2013). Song type matching is an honest early threat signal in a hierarchical animal communication system. Proceedings of the Royal Society of London, Series B: Biological Sciences, 280, 20122517. doi: http://dx.doi.org/10.1098/rspb.2012.2517

Akçay, Ç., Wood, W. E., Searcy, W. A., Templeton, C. N., Campbell, S. E., & Beecher, M. D. (2009). Good neighbour, bad neighbour: song sparrows retaliate against aggressive rivals. Animal Behaviour, 78(1), 97–102.

Anderson, R. C., Searcy, W. A., & Nowicki, S. (2005). Partial song matching in an eastern population of song sparrows, *Melospiza melodia*. Animal Behavior, 69, 189–196.

Anderson, R. C., Searcy, W. A., Peters, S., & Nowicki, S. (2008). Soft song in song sparrows: Acoustic structure and implications for signal function. Ethology, 114(7), 662–676.

Beecher, M. D. (2008). Function and mechanisms of song learning in song sparrows. Advances in the study of behavior, 38, 167–225.

Beecher, M. D., Campbell, S. E., & Burt, J. M. (1994). Song perception in the song sparrow: birds classify by song type but not by singer. Animal Behaviour, 47, 1343–1351.

Beecher, M. D., Campbell, S. E., Burt, J. M., Hill, C. E., & Nordby, J. C. (2000). Song type matching between neighboring song sparrows. Animal Behavior, 59, 21–27.

Beecher, M. D., Campbell, S. E., & Stoddard, P. K. (1994). Correlation of song learning and territory establishment strategies in the song sparrow. Proceedings of the National Academy of Sciences, USA, 91, 1450–1454.

Brainard, M. S., & Doupe, A. J. (2000). Auditory feedback in learning and maintenance of vocal behaviour. Nature Reviews Neuroscience, 1(1), 31–40. doi: 10.1038/35036205

Branch, C. L., Kozlovsky, D. Y., & Pravosudov, V. V. (2015). Elevation related variation in aggressive response to mirror image in mountain chickadees. Behaviour, 152(5), 667–676.

Brooks, R. J., & Falls, J. B. (1975). Individual recognition by song in white-throated sparrows. I. Discrimination of songs of neighbors and strangers. Canadian Journal of Zoology, 53(7), 879–888.

Buniyaadi, A., Taufique, S. K.T., & Kumar, V. (2019). Self-recognition in corvids: evidence from the mirror-mark test in Indian house crows (Corvus splendens). Journal of Ornithology. doi: 10.1007/s10336-019-01730-2

Catchpole, C. K., & Slater, P. J. B. (2008). Bird song: biological themes and variations. Cambridge, UK: Cambridge University Press.

Censky, E. J., & Ficken, M. S. (1982). Responses of black-capped chickadees to mirrors. The Wilson Bulletin, 94(4), 590–593.

Clary, D., & Kelly, D. M. (2016). Graded Mirror Self-Recognition by Clark’s Nutcrackers. Scientific Reports, 6(1), 36459. doi: 10.1038/srep36459

Clary, D., Stow, M. K., Vernouillet, A., & Kelly, D. M. (2020). Mirror-mediated responses of California scrub jays (Aphelocoma californica) during a caching task and the mark test. Ethology, 126(2), 140–152. doi: 10.1111/eth.12954

Derégnaucourt, S., & Bovet, D. (2016). The perception of self in birds. Neuroscience & Biobehavioral Reviews, 69, 1–14.

Falls, J. B. (1985). Song matching in western meadowlarks. Canadian Journal of Zoology, 63(11), 2520–2524.

Falls, J. B., Krebs, J. R., & McGregor, P. (1982). Song matching in the great tit (*Parus major*): The effect of similarity and familiarity. Animal Behaviour, 30(4), 997–1009.

Fisher, J. B. (1954). Evolution and bird sociality. In J. Huxley, A. C. Hardy & E. B. Ford (Eds.), Evolution as process (pp. 71–83). London: Allen & Unwin.

Hill, C. E., Campbell, S. E., Nordby, J. C., Burt, J. M., & Beecher, M. D. (1999). Song sharing in two populations of song sparrows. Behavioral Ecology & Sociobiology, 46, 341–349.

Konishi, M. (1965). The Role of Auditory Feedback in the Control of Vocalization in the White◻Crowned Sparrow 1. Zeitschrift für Tierpsychologie, 22(7), 770–783.

Kraft, F.-L., Forštová, T., Utku Urhan, A., Exnerová, A., & Brodin, A. (2017). No evidence for self-recognition in a small passerine, the great tit (Parus major) judged from the mark/mirror test. Animal Cognition, 20(6), 1049–1057. doi: 10.1007/s10071-017-1121-7

Kroodsma, D. E. (1976). The effect of large song repertoires on neighbor “recognition” in male song sparrows. Condor, 78, 97–99.

Kusayama, T., Bischof, H.-J., & Watanabe, S. (2000). Responses to mirror-image stimulation in jungle crows (Corvus macrorhynchos). Animal Cognition, 3(1), 61–64.

Lawrence, M. A. (2016). Package ‘ez’. R package version, 4.4-0.

Lombardino, A.-J., & Nottebohm, F. (2000). Age at deafening affects the stability of learned song in adult male zebra finches. Journal of Neuroscience.

McArthur, P. D. (1986). Similarity of playback songs to self song as a determinant of response strength in song sparrows (*Melospiza melodia*). Animal Behaviour, 34(1), 199–207.

McArthur, P. D. (1987). Auditory self-perception: A reply to Suarez and Gallup. Anim. Behav., 35(2), 612–613.

McGregor, P. K. (2000). Playback experiments: design and analysis. acta ethologica, 3(1), 3–8. doi: 10.1007/s102110000023

Medina, F., Taylor, A., Hunt, G., & Gray, R. D. (2011). New Caledonian crows’ responses to mirrors. Animal Behaviour, 82(5), 981–993.

Nakagawa, S., & Schielzeth, H. (2010). Repeatability for Gaussian and nonLGaussian data: a practical guide for biologists. Biological Reviews, 85(4), 935–956.

Nice, M. M. (1943). Studies in the life history of the song sparrow II. The behavior of the song sparrow and other passerines. Transactions of the Linnean Society of New York, 6, 1–328.

Nordby, J. C., Campbell, S. E., & Beecher, M. D. (1999). Ecological correlates of song learning in song sparrows. Behavioral Ecology, 10, 287–297.

Nordeen, K. W., & Nordeen, E. J. (1992). Auditory feedback is necessary for the maintenance of stereotyped song in adult zebra finches. Behavioral and neural biology, 57(1), 58–66.

Payne, R. B. (1982). Ecological consequences of song matching: breeding success and intraspecific song mimicry in indigo buntings. Ecology, 63(2), 401–411.

Poirier, C., Boumans, T., Verhoye, M., Balthazart, J., & Van der Linden, A. (2009). Own-song recognition in the songbird auditory pathway: selectivity and lateralization. Journal of Neuroscience, 29(7), 2252–2258.

Prather, J. F., & Mooney, R. (2004). Neural correlates of learned song in the avian forebrain: simultaneous representation of self and others. Current Opinion in Neurobiology, 14(4), 496–502.

Price, P. H. (1979). Developmental determinants of structure in zebra finch song. Journal of Comparative and Physiological Psychology, 93, 260–277.

Prior, H., Schwarz, A., & Güntürkün, O. (2008). Mirror-induced behavior in the magpie (Pica pica): evidence of self-recognition. PLoS biology, 6(8), e202.

R Core Team. (2012). R: A language and environment for statistical computing. Vienna, Austria: R Foundation for Statistical Computing.

Searcy, W. A., Anderson, R. C., & Nowicki, S. (2006). Bird song as a signal of aggressive intent. Behavioral Ecology and Sociobiology, 60(2), 234–241.

Searcy, W. A., McArthur, P. D., Peters, S. S., & Marler, P. (1981). Response of Male Song and Swamp Sparrows to Neighbour, Stranger, and Self Songs. Behaviour, 77, 152–163.

Searcy, W. A., Ocampo, D., & Nowicki, S. (2019). Constraints on song type matching in a songbird. Behavioral Ecology and Sociobiology, 73(8), 102.

Soler, M., Pérez-Contreras, T., & Peralta-Sánchez, J. M. (2014). Mirror-Mark Tests Performed on Jackdaws Reveal Potential Methodological Problems in the Use of Stickers in Avian Mark-Test Studies. PLOS ONE, 9(1), e86193. doi: 10.1371/journal.pone.0086193

Stoddard, P. K., Beecher, M. D., Horning, C. H., & Campbell, S. E. (1992). Song type matching in the song sparrow. Canadian Journal of Zoology, 70, 1440–1444.

Stoddard, P. K., Beecher, M. D., Horning, C. L., & Campbell, S. E. (1991). Recognition of individual neighbors by song in the song sparrow, a species with song repertoires. Behavioral Ecology and Sociobiology, 29(3), 211–215.

Suarez, S. D., & Gallup, G. G. (1987). The question of an auditory self-concept in song sparrows, Melospiza melodia.

Temeles, E. J. (1994). The role of neighbors in territorial systems: when are they ‘dear enemies’? Animal Behavior, 47, 339–350.

Vanhooland, L.-C., Bugnyar, T., & Massen, J. J. (2019). Crows (Corvus corone ssp.) check contingency in a mirror yet fail the mirror-mark test. Journal of Comparative Psychology.

Weeden, J. S., & Falls, J. B. (1959). Differential responses of male ovenbirds to recorded songs of neighboring and more distant individuals. Auk, 76, 343–351.

Yasukawa, K., Bick, E., Wagman, D., & Marler, P. (1982). Playback and speaker-replacement experiments on song-based neighbor, stranger, and self discrimination in male red-winged blackbirds. Behavioral Ecology and Sociobiology, 10(3), 211–215.

